# CIC is a Critical Regulator of Neuronal Differentiation

**DOI:** 10.1101/799536

**Authors:** Inah Hwang, Heng Pan, Jun Yao, Olivier Elemento, Hongwu Zheng, Jihye Paik

## Abstract

Capicua (CIC), a member of the high mobility group (HMG)-box superfamily of transcriptional repressors, is frequently mutated in human oligodendrogliomas. But its function in brain development and tumorigenesis remains poorly understood. Here, we report that brain-specific deletion of *Cic* compromises developmental transition of neuroblast to immature neurons in mouse hippocampus and compromises normal neuronal differentiation. Combined gene expression and ChIP-seq analyses identified VGF as an important CIC-repressed transcriptional surrogates involved in neuronal lineage regulation. Aberrant VGF expression promotes neural progenitor cell proliferation by suppressing their differentiation. Mechanistically, we demonstrated that CIC represses VGF expression by tethering SIN3-HDAC to form a transcriptional corepressor complex. Mass spectrometry analysis of CIC-interacting proteins further identified BRG1 containing mSWI/SNF complex of which function is necessary for transcriptional repression by CIC. Together, this study uncovers a novel regulatory pathway of CIC-dependent neuronal differentiation and provides molecular insights into the etiology of CIC-dependent brain tumors.

## Introduction

Oligodendroglioma, a diffusely infiltrating primary malignant brain tumor of adult, is histologically characterized by its composition of neoplastic cells morphologically resembling oligodendroglial cells. Despite of their generally better prognosis and response to early chemotherapy, most oligodendroglial tumors recur eventually, with many of them progressing to higher grade lesion (Cairncross, Wang et al., 2013). Genetically, oligodendrogliomas are defined by a combined loss of chromosome arms 1p and 19q (Cairncross, Ueki et al., 1998). More recent high-throughput sequencing approach has further identified *CIC* (*homolog of Drosophila capicua*) gene, which is mapped to 19q13.2, as one of the most frequently mutated genes in oligodendrogliomas (Bettegowda, Agrawal et al., 2011, Jiao, Killela et al., 2012, Yip, Butterfield et al., 2012), pointing to a critical role of CIC in brain development and oligodendroglioma pathogenesis.

CIC is a transcriptional repressor with a SOX-like high mobility group (HMG) DNA binding domain (Lee, Chan et al., 2002). Studies in *Drosophila* revealed CIC as a key mediator of RAS/MAPK signaling that regulates embryonic patterning and intestinal stem cell proliferation (Jiménez, Guichet et al., 2000, Jin, Ha et al., 2015). In mammals, CIC has been shown to play a critical role in T cell development and adaptive immunity (Park, Lee et al., 2017, Tan, Brunetti et al., 2018), lung alveolarization and abdominal wall closure during the development (Lee, Fryer et al., 2011, Simón-Carrasco, Graña et al., 2017), and bile acid homeostasis (Kim, Park et al., 2015). As to its function in central nervous system (CNS) development, interestingly, a previous study of mouse brain-specific Cic deletion by glial fibrillary acidic protein promoter driven cre recombinase (hGFAP-cre) reported no gross developmental abnormalities (Simón-Carrasco et al., 2017). By contrast, another study using *Emx1-cre* revealed that CIC was important for neuro-development (Lu, Tan et al., 2017). Loss of CIC disrupted organization and maintenance of upper layer cortical neurons and led to mouse hyperactivity with defective learning and memory loss. More recently, an additional study using also *Foxg1-cre* revealed that mouse forebrain-specific deletion of *Cic* caused abnormal increase of oligodendrocyte progenitor cells (OPC) and immature oligodendrocytes populations (Ahmad, Rogers et al., 2019, Yang, Chen et al., 2017), likely at the expense of neuronal propagation. But despite those efforts, the molecular function(s) of CIC in brain development and tumorigenesis still remains poorly understood. Here, we describe a new mouse model in which we applied embryonic neural progenitor cells (NPC) targeting Nestin-cre (*Nes-cre*) to broadly inactivate CIC in CNS. Our data show that the loss of CIC compromises the normal developmental transition of neuroblasts to neurons and therefore their differentiation. Through integrated expression and ChIP-seq analyses, we identified VGF as a novel CIC surrogates that mediates its function in regulating neuronal lineage differentiation. We further show that CIC transcriptionally represses its target gene expression by recruiting mSWI/SNF and SIN3-HDAC repressor complexes. Our findings suggest CIC as a critical developmental transcriptional repressor that blocks tumorigenesis by facilitating neuronal maturation.

## Results

### Defective cerebral cortex development in *Cic^KO^* mouse

To investigate the role of CIC in neurogenesis, we engineered a conditional *Cic* floxed mouse allele and crossed it with *Nes-cre* animal to target *Cic* deletion in embryonic neural precursors and their progenies (Fig EV1A). Recombination of floxed *Cic* allele in brains of *Nes-cre; Cic^Lox/lox^* (*Cic^KO^*) animals was confirmed by PCR (Fig EV1B-D). Developmentally, the *Cic^KO^* mice displayed severe growth retardation and reduction of brain volume by postnatal day 14 (Fig 1A and 1B). Although largely indistinguishable from their littermate wild-type and heterozygous counterparts at birth, the growth retardation became evident in *Cic^KO^* pups by postnatal day 6. None of *Cic^KO^* pups survived beyond postnatal day 20-22 (N = 21). The efficiency of CIC depletion was confirmed by immunofluorescence (IF) analysis (Fig 1C). Histologic examination of *Cic^KO^* pups at postnatal day 14 further revealed significant reduction of cerebral cortex thickness compared to *Cic^WT^* littermates (N = 10) (Fig 1D), indicating that CIC is required for early brain development.

**Figure 1.**
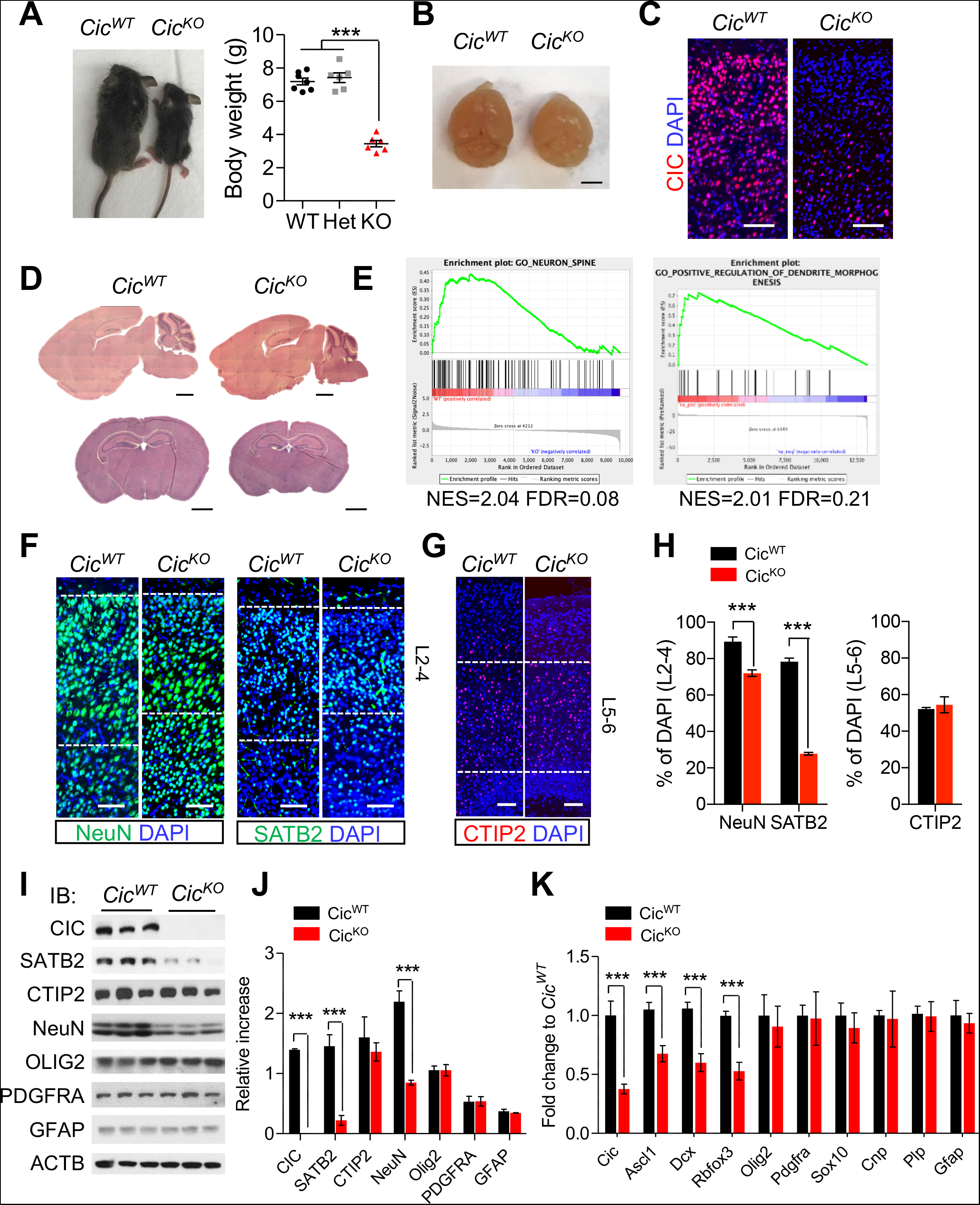
Defective cerebral cortex developments in *Cic^KO^* mouse. A. The picture (left) and the plot for body weights (right) at postnatal 14 days. B. The brains at postnatal 14 days. C. IF analysis for CIC in the cerebral cortex, scale bar: 100 μm. D. H&E staining of *Cic^WT^* and *Cic^KO^* brains, scale bar: 1 mm. E. GSEA in *Cic^KO^ vs Cic^WT^* brains. F, G. IF analysis for NeuN, SATB2, and CTIP2 in the cerebral cortex of P14 mouse, scale bar: 100 μm. H. Quantifications at layer 2-4 of NeuN or SATB2 positive (G) and layer 5-6 of CTIP2 positive (H) numbers are plotted. Mean ± SEM of 200 DAPI positive nuclei from 3 animals, ***p<0.001 (Unpaired t-test). I. WB analysis for indicated protein expressions in cerebral cortex lysates of P14 *Cic^WT^* and *Cic^KO^* brains. J. Densitometry analysis of multiple WB in (J) is plotted. Mean ± SEM of 3 experimental animals, ***p<0.001 (Unpaired t-test). K. qPCR results of indicated genes in P14 *Cic^KO^ vs Cic^WT^* brains. Mean ± SEM of 6 experimental animals, ***p<0.001 (Unpaired t-test).

To uncover the underlying cause of the observed brain phenotype, we performed RNA sequencing (RNA-seq) analysis of P0.5 brains from control and *Cic^KO^* animals. Gene set enrichment analysis (GSEA) (Mootha, Lindgren et al., 2003, Subramanian, Tamayo et al., 2005) revealed that downregulated genes in *Cic*^KO^ mouse were enriched in the gene sets related with neuronal development (Fig 1E). Indeed, IF analysis of P14 animals showed markedly reduced numbers of NeuN-positive neurons in cortical layer 2-4 compared to those of *Cic^WT^* controls. Interestingly, although the number of SATB2-positive cortical neurons in the layer 2-4 was evidently reduced in P14 *Cic^KO^* animals, we did not find significant differences in the number of CTIP2-positive cortical neurons in layer 5-6 when compare to those of P14 *Cic^WT^* control pups (Fig 1F-H). These observations were further corroborated by both western blot (WB) and quantitative real-time PCR (qPCR) analysis of brain cortices from corresponding animals (Fig 1I-K), suggesting that CIC is necessary for the normal development of the cerebral cortex.

Notably, two previous studies (Ahmad et al., 2019, Yang et al., 2017) reported that deletion of *Cic* caused increased number of either OLIG2/PDFGRA-positive OPC or GFAP-positive astrocytes in the affected mouse brain. To test this in our setting, we next analyzed the expression of various lineage specific genes for oligodendrocytes (*e.g., Olig2, Pdgfra, Sox10, Cnp*, and *Plp*), astrocytes (*e.g., Gfap*) and neurons (*e.g., Ascl1, Dcx*, and *Rbfox3*) in brain samples derived from P14 *Cic^KO^* and their littermate *Cic^WT^* animals. To our surprise, although the expression of neuronal genes was consistently reduced in *Cic^KO^* compared to their *Cic^WT^* control animals, there was no evident differences in either oligodendrocytic or astrocytic lineage marker expressions (Fig 1K). WB and GSEA analysis of P0.5 animals confirmed that neither downregulated nor upregulated genes in *Cic*^KO^ mouse were significantly enriched in gene sets related with oligodendrocyte or astrocyte differentiation (Fig EV2A-C). Consistently, the immunohistochemistry (IHC) analysis did not show abnormal expansion of OLIG2-or GFAP-positive cells in P14 *Cic^KO^* brains (Fig EV2D). In addition, we did not observe reduction of myelin basic protein (MBP) staining in P14 *Cic^KO^* brains (Fig EV2D), suggesting CIC is mainly involved in neuronal lineage differentiation regulation during brain development.

### Loss of CIC compromises neuronal maturation

Next, we determined CIC protein expression in different CNS cell types to understand its role in brain development. IF analysis of early postnatal and adult mouse brains revealed that CIC is highly expressed in NeuN-positive neurons (Fig 2A-B). But its expression in GFAP-positive astrocytes and OLIG2-positive oligodendrocytes is much weaker than that of neurons (Fig EV3A-D). This is consistent with the finding that CIC is primarily involved in regulation of neurogenesis.

**Figure 2.**
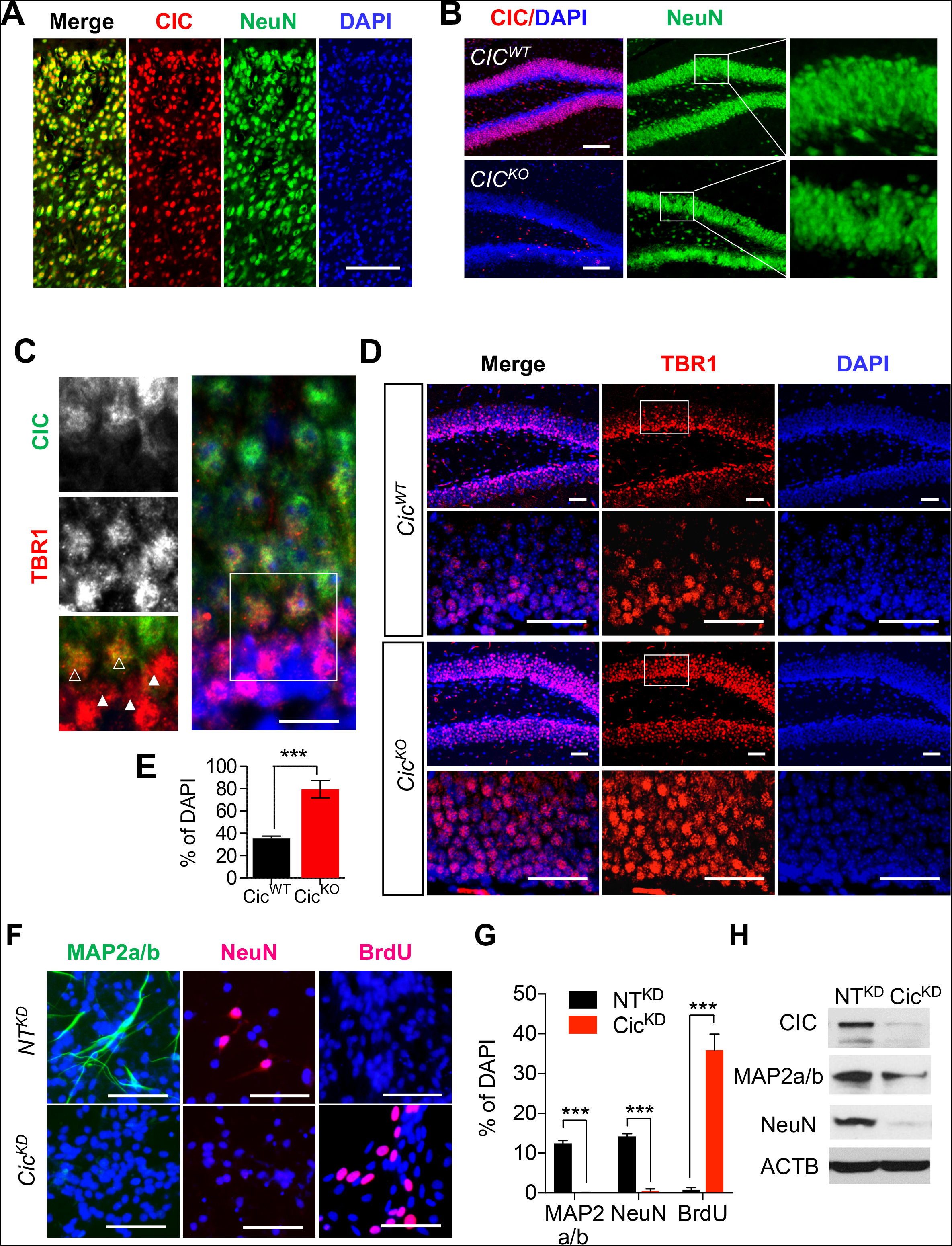
Loss of CIC compromises neuronal maturation. A, B. IF analysis for CIC and NeuN, scale bar: 200 μm. C. Co-IF for TBR1 and CIC expression in hippocampus. Open arrow heads point to TBR1-low cells and closed ones are on TBR1-high cells. D. IF analysis for TBR1 in P14 *Cic^WT^* and *Cic^KO^* brains, scale bar: 50 μm. E. TBR1 high cells of (D) images are counted. Mean ± SEM of the measurement from 50-60 DAPI from 3 animals, ***p<0.001 (Unpaired t-test). F. IF analysis for MAP2a/b, NeuN, and BrdU in NT^KD^ NPC and Cic^KD^ NPC at 8 days of differentiation, scale bar: 50 μm. G. Quantitation of experiments in (F). Mean ± SEM of the measurement from 100 DAPI positive nuclei from 3 experiments, *** p<0.001(Unpaired t-test). H. WB analysis for each specific marker at 5 days of differentiation.

To examine a potential function of CIC along the neuronal differentiation process, we measured cellular CIC expression in the neurogenic subgranular zone (SGZ) and subventricular zone (SVZ) areas of adult mouse brains. Co-immunostaining with a panel of stage specific markers showed a pattern of progressively increased CIC expression along the neuronal differentiation. Specifically, compared to the mature NeuN-positive neurons that ubiquitously registered high level of CIC expression (Fig 2A and 2B), the Nestin (or SOX2)-positive type 1 (or type B) neural stem cells, TBR2-postive type 2A, ASCL1-postive type C transit-amplifying progenitor cells, or DCX-positive neuroblasts all exhibited relatively low CIC expression (Fig EV4A-C). Consistently, CIC expression was weak in TBR1-high neuroblast cells or immature neurons but became increasingly strong later in TBR1-low or NeuN-positive mature neurons (Fig 2C). This pattern of dynamically regulated CIC expression along the neurogenesis process supports the premise that it plays an important role during neuronal maturation. Indeed, the analysis of hippocampal granular layers of *Cic^KO^* brains revealed abnormally expanded TBR1-high cells but reduced populations of TBR1-low cells compared to the *Cic^WT^* controls (Fig 2D and 2E). As the consequence, the density of dentate gyrus NeuN+ granular layer was also noticeably decreased in the *Cic^KO^* animals relative to that of the littermate controls (Fig 2B).

To further explore the function of CIC in neuronal differentiation, we adopted a previously established *in vitro* differentiation system by introducing tet-inducible ASCL1 into immortalized neural progenitors (NPC) (Hwang, Cao et al., 2018). Sixty percent of NPC highly expressed ASCL1 and were rapidly differentiated to DCX-positive neuroblast upon treatment of doxycycline (Fig EV5A and EV5B). Similar to primary neural stem/progenitor cells, these cells could be further differentiated to NeuN positive neurons (Fig EV5C and EV5D) and cease to proliferate as evidenced by lack of BrdU incorporation (Fig EV5E). To determine the role of CIC in neuronal differentiation, we generated *Cic*-knocked-down and control NPC using short hairpin RNA (Cic^KD^ and NT^KD^). Importantly, NeuN-positive mature neurons were not detected in Cic^KD^ NPC on day 8 of differentiation unlike NT^KD^ NPC (Fig 2F-H). To further determine whether the loss of CIC sustains proliferation, we analyzed differentiated cells for BrdU incorporation. On day 8 of differentiation, ∼20% of Cic^KD^ NPC continued to be BrdU-positive (Fig 2F-H). These results suggest that CIC is necessary for complete exit from cell cycle and terminal neuronal differentiation.

### CIC represses transcriptional targets in developing brains

CIC is a member of the high mobility group (HMG)-box superfamily of transcriptional repressor (Lee et al., 2002). To identify its direct transcriptional surrogates that may play a role in neurogenesis, we performed chromatin immunoprecipitation followed by DNA sequencing (ChIP-Seq) concurrently with RNA-seq using P0.5 control and *Cic^KO^* mouse brains that are undergoing neuronal maturation. Two biological replicates showed concordant genome-wide peaks that are absent in input control (Fig 3A), verifying quality of the data. *De novo* motif analysis further revealed that CIC bindings are strongly enriched at the AT-rich regions (Fig 3B). A significant portion of CIC peaks were resided within the promoter regions (∼ 5%) or introns (∼ 20%), whereas the rest were in intergenic regions (∼ 45%). In agreement with its presumed role as a transcriptional repressor, RNA-seq comparisons of P0.5 *Cic^WT^* control and *Cic^KO^* brain samples revealed that CIC depletion increased the expression of genes with CIC interactive promoters. Out of 50 most differentially upregulated, 13 genes have CIC peaks in their promoter regions suggesting direct transcriptional repression by CIC binding (Fig 3C and table EV1). Alignment of CIC peaks with corresponding H3K27Ac ChIP-seq annotations confirmed that signal density of active promoter/enhancer-designated H3K27Ac peak centers was higher in *Cic^KO^* samples than that of *Cic^WT^* (Fig 3D). By contrast, the expression of genes with only intronic CIC peaks (*e.g., Skap2, Lipa, and Cntnap2*) did not show significant changes following CIC loss, suggesting that CIC may exert its transcriptional repression function mainly at promoter/enhancer regions.

**Figure 3.**
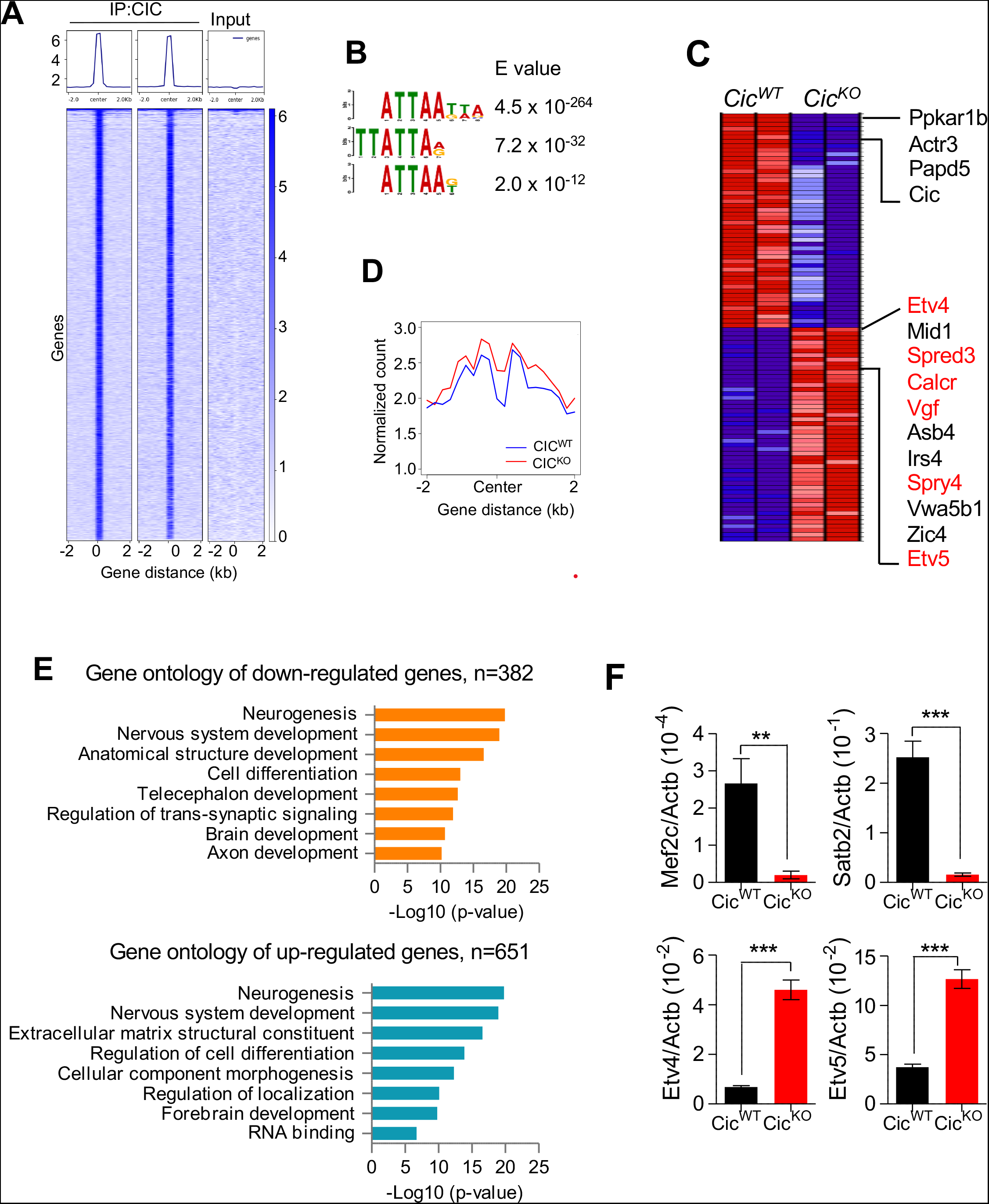
CIC represses transcriptional targets in developing brains. A. Heatmaps at CIC peaks showing CIC and input. B. Motif analysis. C. Heatmap of the top 50 ranked differentially expressed genes in RNA-seq analysis of P0.5 brains of *Cic^KO^* vs. *Cic^WT^*. Genes with CIC ChIP peaks are marked red. D. Average ChIP-seq signal of H3K27Ac at CIC peaks from P0.5 brains of *Cic^KO^* vs. *Cic^WT^*. E. Gene ontology analysis with RNA-seq in P0.5 brains of *Cic^KO^* vs. *Cic^WT^*. The up-or down-regulated genes were retrieved with p<0.02 (Unpaired t-test). F. qPCR results of *Mef2c, Satb2, Etv4*, and *Etv5* mRNA expressions in P0.5 brains of *Cic^WT^* and *Cic^KO^*. Mean ± SEM of 5 experimental animals, **p<0.01, *** p<0.001 (Unpaired t-test).

To explore underlying cellular and developmental mechanisms associated with CIC loss, we next performed the gene ontology comparison of gene expression of *Cic^WT^* control and *Cic^KO^* brain samples. Annotation of the 1033 differentially expressed genes (p<0.02) indeed revealed neurogenesis as the mostly dysregulated biological process following CIC depletion (Fig 3E), which is consistent with phenotype observed in the *Cic^KO^* brains. qPCR analysis further confirmed that expression of the neural transcriptional regulators *Mef2c* and *Satb2* were significantly downregulated in *Cic^KO^* brains (Fig 3F). In contrast, previously reported CIC transcriptional targets (*e.g., Etv4, Etv5*) were upregulated as in *Cic^KO^* brains.

### Aberrant expression of VGF precludes development of mature neurons in CIC deficient cells

We next leveraged transcriptional and ChIP-seq profiles of the paired *Cic^WT^* control and *Cic^KO^* brains to identify novel CIC transcriptional targets that are potentially involved in regulation of neurogenesis. Among those exhibited strong CIC binding in their promoter regions, the mRNA expression of *Vgf* (VGF nerve growth factor inducible) was consistently upregulated in *Cic^KO^* brains (Fig 3C and 4A). VGF is a neuropeptide that is important not only for neuronal maturation during development (Alder, Thakker-Varia et al., 2003), but also plays a crucial role promoting glioma stem cell survival and stemness in glioma pathogenesis (Wang, Prager et al., 2018). Consistent with its increased mRNA expression following CIC depletion, WB and IF staining showed that protein levels of VGF were markedly elevated in *Cic^KO^* brain lysates and cortex in comparison to those of wildtype counterpart (Fig 4B and 4C). Increased VGF protein expression was also observed in cultured NPC and their in vitro differentiated progenies following *Cic* knockdown (Fig 4D). Importantly, enforced VGF expression compromised neuronal differentiation capacity of NPC and blocked its cell cycle exit under differentiation induction (Fig 4E-H), recapitulating the phenotype observed in *Cic^KD^* NPC. These findings establish VGF as a crucial mediator of CIC function during neurogenesis.

**Figure 4.**
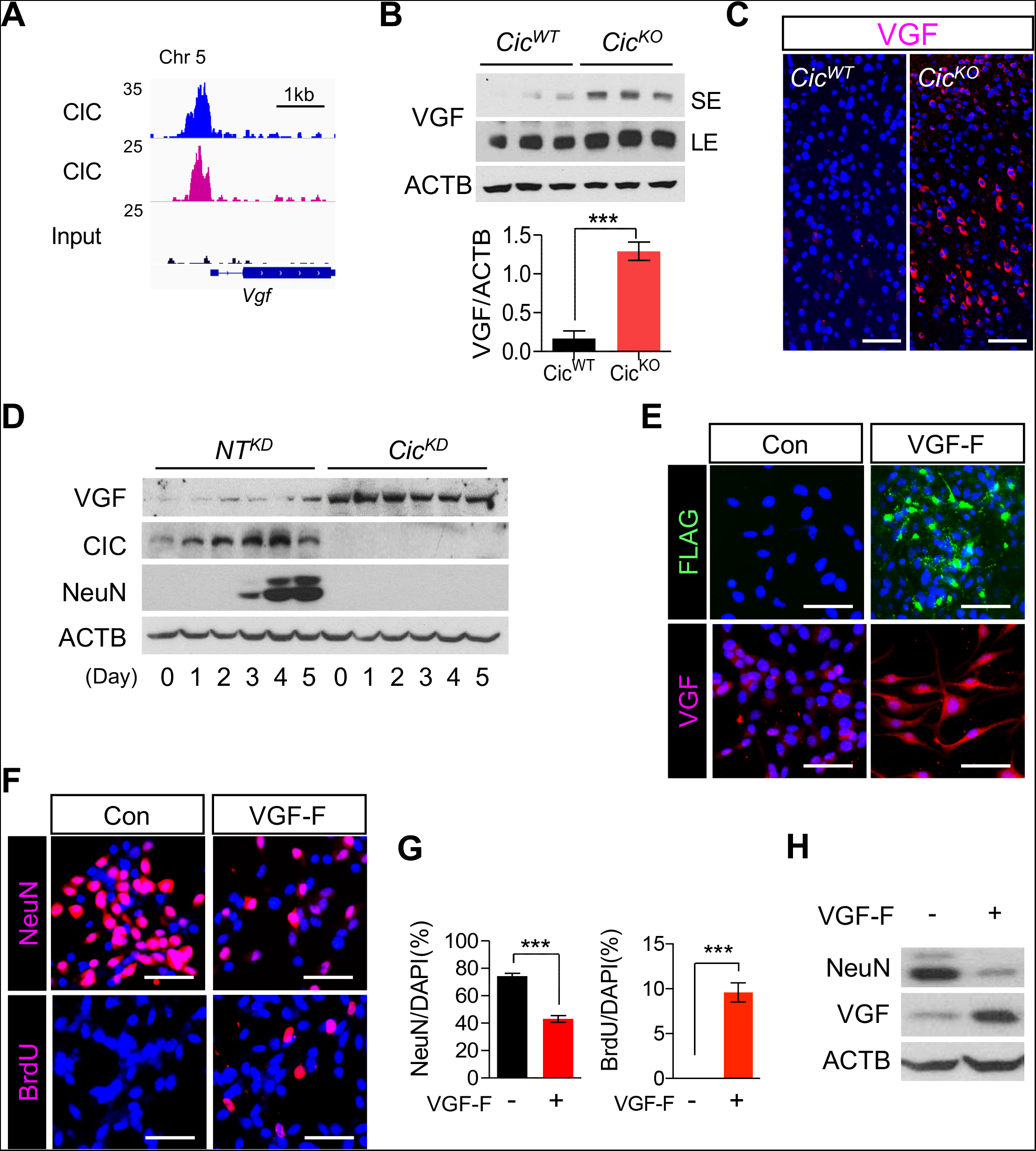
Aberrant expression of VGF compromises neurogenesis in CIC-deficient brains. A. Biological replicates of CIC ChIP-seq tracks at *Vgf* promoter regions. B. WB analysis for VGF in P0.5 brains of *Cic^WT^* and *Cic^KO^*. SE: Short exposure, LE: Long exposure. Mean ± SEM of 3 experimental animals, ***p<0.001 (Unpaired t-test). C. IF analysis for VGF in the P14 brains of *Cic^WT^* and *Cic^KO^*, scale bar: 100 μm. D. WB analysis for VGF in NT^KD^ NPC and Cic^KD^ NPC during differentiation. E. IF analysis for Flag and VGF in VGF-Flag-expressing NPC (VGF-F), scale bar: 50 μm. F. IF analysis for NeuN and BrdU at 7 days of differentiation, scale bar: 50 μm. G. Quantitation for NeuN or BrdU positive numbers (F) are plotted. Mean ± SEM of 100 DAPI positive nuclei from 3 experiments, *** p<0.001 (Unpaired t-test). H. WB analysis for indicated proteins in control NPC and VGF-F expressing NPC at 5 days of differentiation.

### CIC transcriptionally represses VGF expression

To confirm VGF as a direct CIC transcriptional target, we analyzed ChIP-seq profile from paired P0.5 *Cic^WT^* and *Cic^KO^* brain samples. The CIC ChIP-seq track revealed an evident peak at the vicinity of the transcription start site (TSS) of *Vgf* gene (Fig 4A and 5A). The *Vgf* promoter-specific CIC binding was verified by ChIP-qPCR results from two independent sets P0.5 brain samples (Fig 5B). In agreement with its function as a member of high HMG-box superfamily of transcriptional repressors, ChIP-seq and qPCR analysis from independent sets of paired *Cic^WT^* and *Cic^KO^* brain samples showed that brain-specific CIC depletion elicited significant increase of H3K27 acetylation within the *Vgf* promoter region and also elevation of *Vgf* mRNA expression (Fig 5A-C). Similar enrichment of *Vgf* promoter-specific CIC occupancy, regional increase of H3K27Ac density, and correlated increase in its mRNA level following CIC depletion were also observed in cultured NPC and their differentiated progenies (Fig 5D and 5E). Together, these findings indicate that VGF is a bona fide CIC transcriptional downstream target.

**Figure 5.**
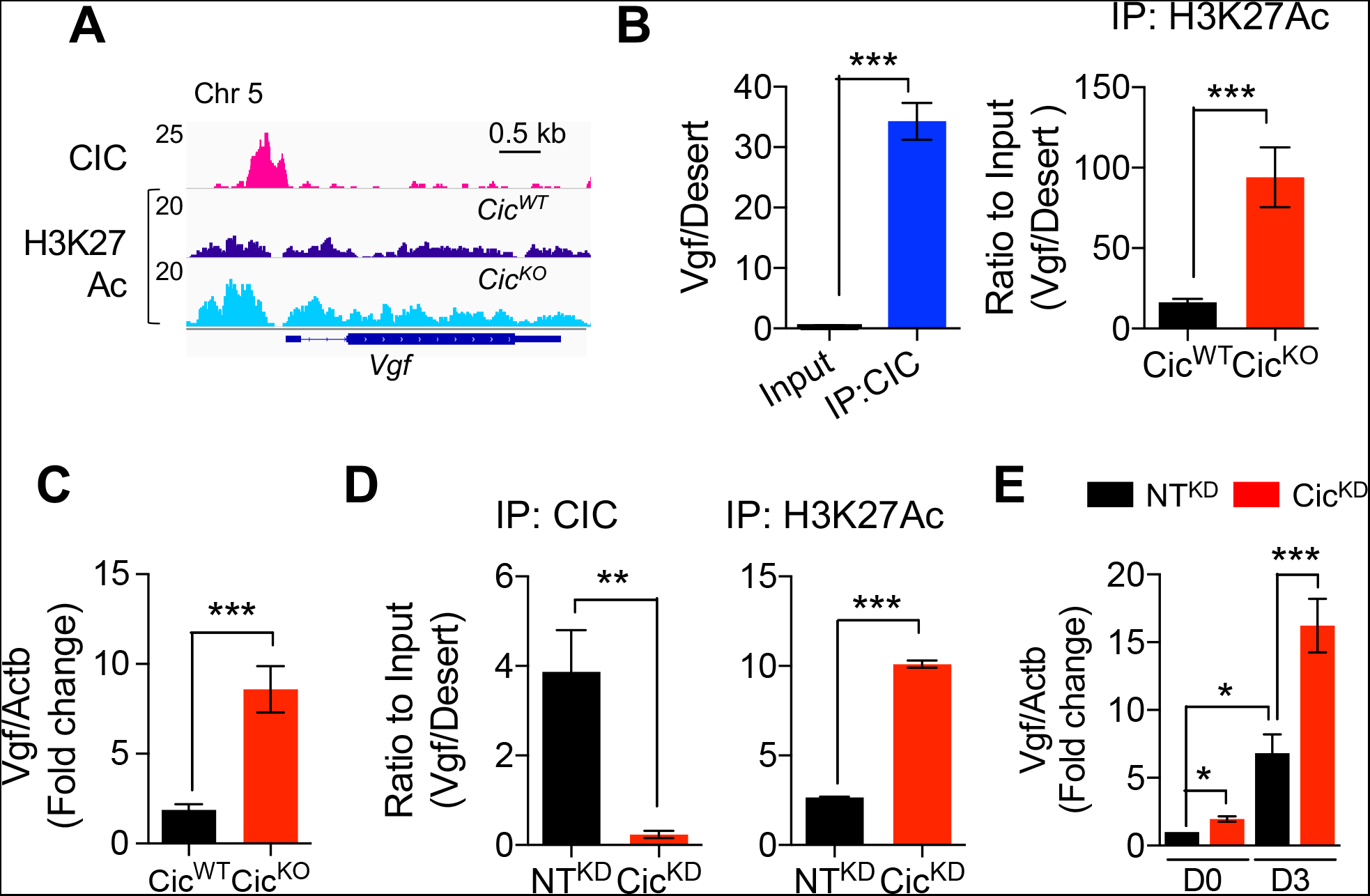
CIC transcriptionally represses VGF. A. Indicated ChIP-seq tracks at *Vgf* promoter region. B. ChIP-qPCR results from P0.5 brains at *Vgf* promoter region. Mean ± SEM of 3 experimental animals, *** p<0.001 (Unpaired t-test). C. qPCR results of *Vgf* mRNA expressions in P0.5 cerebral cortex of *Cic^WT^* and *Cic^KO^*. Mean ± SEM of 8 experimental animals, *** p<0.001 (Unpaired t-test). D. ChIP-qPCR results from NT^KD^ NPC and Cic^KD^ NPC at *Vgf* promoter region. Mean ± SEM of 3 experiments, *** p<0.001 (Unpaired t-test). E. qPCR results of *Vgf* mRNA expression in the NT^KD^ NPC and Cic^KD^ NPC at day 0 (D0) and day 3 (D3) of differentiation. Mean ± SEM of 3 experiments, *p<0.05, *** p<0.001 (One-way ANOVA).

### CIC interacts with mSWI/SNF complex during neurogenesis

To explore the molecular mechanism underlying CIC-mediated transcriptional repression during neurogenesis, we performed CIC immunoprecipitation (IP) from P1 brain lysates followed by mass spectrometry (MS). Besides the previously reported CIC-interacting proteins like ACLY (Chittaranjan, Chan et al., 2014) and ATXN1 (Lam, Bowman et al., 2006), the IP-MS analysis uncovered many core components of mSWI/SNF (mammalian Switch/Sucrose Non-Fermentable) complex, including ARID1A, ARID1B, SMARCC1 (BAF155), SMARCC2 (BAF170), SMARCA2 (BAF190B, BRM), and BRG1 (SMARCA4) (Fig 6A and table EV2). The specific interaction of CIC with mSWI/SNF complex was further verified by co-IP analysis of either CIC-transfected HEK293 cells (Fig 6B) or P0.5 *Cic^WT^* mouse brains (Fig 6C).

**Figure 6.**
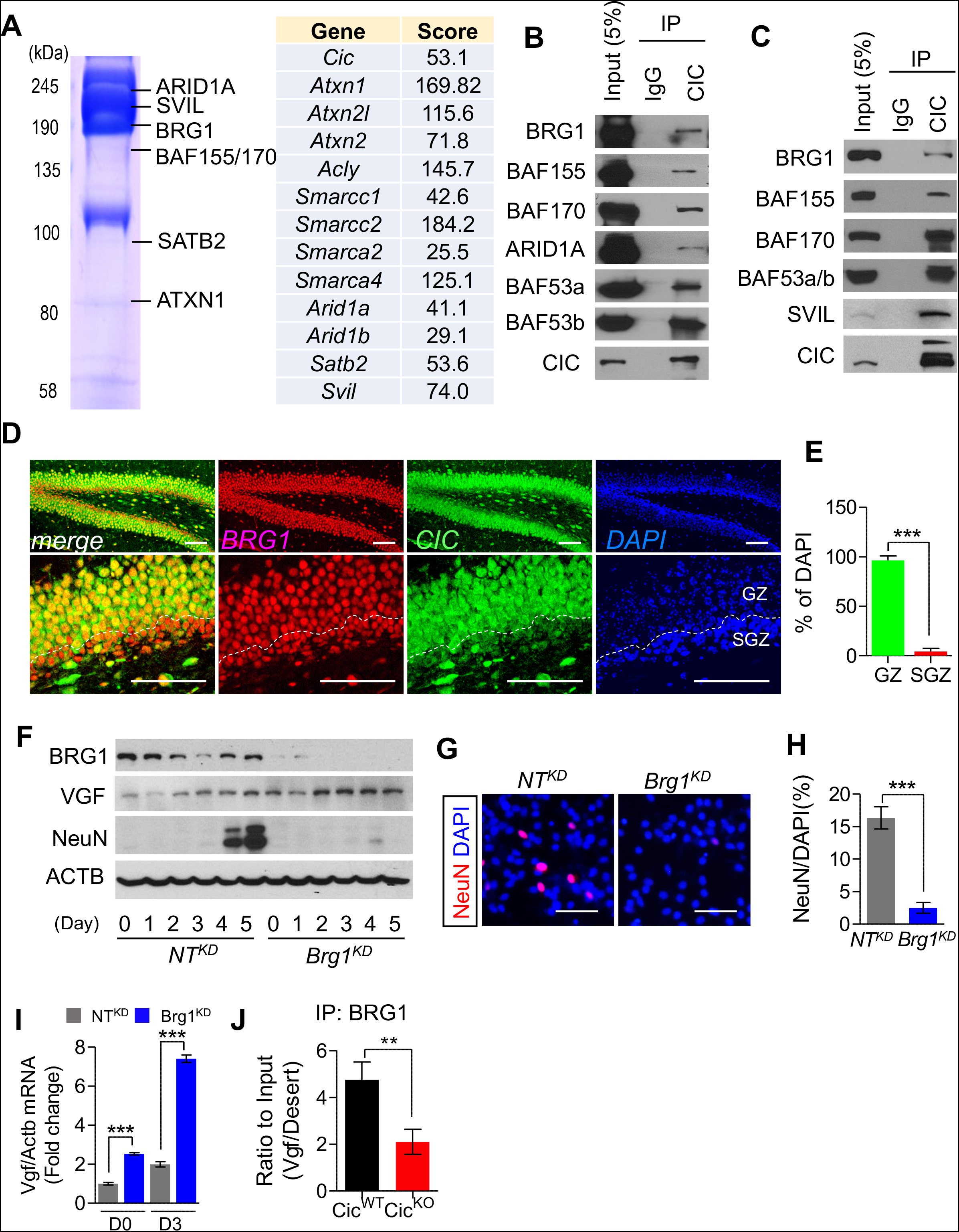
mSWI/SNF complex is a molecular partner for CIC-dependent transcriptional regulation. A. The partial list of CIC interacting proteins analyzed by mass spectrometry. B. Co-IP analysis of CIC with components mSWI/SNF complex after expressing in HEK293T cells. C. Co-IP analysis of endogenous proteins from P0.5 brains. D. Co-IF analysis for CIC and BRG1 in P14 brains, scale bar: 100 μm. E. Ratio of CIC and BRG1 co-expressed cells to DAPI positive nuclei in (D). GZ: granular zone. Mean ± SEM of the measurement from 50 DAPI positive nuclei from 3 animals, *** p<0.001 (Unpaired t-test). F. WB analysis for indicated proteins in NT^KD^ NPC and Brg1^KD^ NPC during differentiation. G. IF analysis for NeuN at 7 days of differentiation, scale bar: 50 μm. H. Quantitation of NeuN positive cells from (G). Mean ± SEM of 100 DAPI positive nuclei from 3 animals, ***p<0.001 (Unpaired t-test). I. qPCR results for *Vgf* mRNA expression in NT^KD^ NPC and Brg1^KD^ NPC at day 0 (D0) and day 3 (D3) of differentiation. Mean ± SEM of 3 experiments, ***p<0.001 (One-way ANOVA). J. ChIP-qPCR results from IP of BRG1 from NT^KD^ NPC and Brg1^KD^ NPC at *Vgf* promoter regions. Mean ± SEM of 3 experiments, **p<0.01 (Unpaired t-test).

The mSWI/SNF, an ATP-dependent multi-subunit chromatin remodeling complex, plays critical roles in the regulation of neural stem cell proliferation, neurogenesis and neocortical development (Ninkovic, Steiner-Mezzadri et al., 2013, Sokpor, Xie et al., 2017, Yu, Chen et al., 2013). Indeed, immunostaining found that BRG1, the core subunit of mSWI/SNF, was highly expressed in the granular cell layer of hippocampus, similar to the CIC expression pattern (Fig 6D-E). To test whether disruption of mSWI/SNF complex would recapitulate CIC loss-associated developmental and molecular phenotypes, we applied CRISPR/Cas9-based genomic editing to generate BRG1 depleted (Brg1^KD^) NPC (Fig 6F). Depletion of BRG1 not only elevated VGF expression in the NPC and its differentiated progenies (Fig 6F), it also compromised their differentiation capacity and neuronal marker expression during neuronally-directed differentiation induction (Fig 6F-H). In addition, BRG1 ChIP-qPCR from P0.5 *Cic^WT^* mouse brains showed enriched occupancy within the *Vgf* promoter region (Fig 6J). Importantly, this BRG1 binding was significantly reduced in *Cic^KO^* brains (Fig 6J), indicating that mSWI/SNF-mediated chromatin remodeling in regulation of VGF transcription requires CIC.

In addition to the mSWI/SNF complex, we also identified SVIL, an actin-binding protein encoded by *supervillin* gene, as a CIC-and SWI/SNF complex-associated protein (Fig 6A and 6C). IP of BRG1 followed by immunoblot analysis of P0.5 brain lysates confirmed the interaction of SVIL with mSWI/SNF complex (Appendix Fig S1A). SVIL has been implicated in the regulation of neuronal differentiation (Laurent, Ruitu et al., 2015). Interestingly, IF analysis of NPC revealed that SVIL was diffusively distributed throughout the cytoplasm; but it became mostly nuclear as the NPC underwent neuronal differentiation induction (Appendix Fig S1B). To search the motifs or amino acid residues critical for its localization switch, we performed MotifScan and identified S243 within its N-terminal nuclear localization signal region as a strong CDK5 phosphorylation site (Appendix Fig S1C). Since CDK5 plays an important role in neuronal differentiation (Jessberger, Gage et al., 2009), we next tested whether its activity is required for differentiation-induced SVIL nuclear translocation. Indeed, treatment of NPC with CDK5 inhibitor roscovitine blocked the nuclear switch of SVIL when the cells were subjected to neuronal differentiation induction and compromised their differentiation (Appendix Fig S1D and S1E). Similarly, knockdown of SVIL (Svil^KD^) in NPC also suppressed their neuronal differentiation capacity as evidenced by the lack of mature neuronal marker MAP2a/b and NeuN expressions. In addition, compared to the control NPC that fully underwent differentiation and exited cell cycle, approximately 5 % of the SVIL-depleted cells continued to incorporate BrdU even after 8 days of differentiation induction (Appendix Fig S1F-H). Together, these findings indicate that SVIL functions together with CIC-mSWI/SNF complex to facilitate neuronal differentiation.

### CIC tethers SIN3-HDAC corepressor complex and mSWI/SNF complex to VGF promoter during neurogenesis

In support of the hypothesis we found that CIC and BRG1 (Attanasio, Nord et al., 2014) showed largely overlapping peaks at the promoter regions of CIC target genes in developing mouse brains (Fig 7A). Consistent with the expression levels, H3K27Ac coverage in CIC peak region was consistently higher on these genes in *Cic^KO^* than *Cic^WT^* brains suggesting HDAC-dependent transcriptional repression might be a mechanism. SIN3 is a scaffold for HDAC-associated transcriptional co-repressor complex (Laherty, Yang et al., 1997). Since SIN3 can interact with CIC and also mSWI/SNF complex through BRG1 (Kuzmichev, Zhang et al., 2002, Weissmann, Cloos et al., 2018), we next tested whether CIC-mSWI/SNF complex-mediated repression of VGF transcription necessitated SIN3 co-repressor complex. ChIP-qPCR analysis of NPC samples confirmed significantly enriched occupancies of SIN3A and neuronal SIN3 co-repressor complex associated HDAC2 within the *Vgf* promoter region where CIC and BRG1 peaks overlap (Baltan, Bachleda et al., 2011) (Fig 7B). Importantly, the regional enrichment of BRG1, SIN3A and HDAC2 were all evidently reduced following CIC depletion, indicating their recruitment to *Vgf* promoter is dependent on the presence of CIC. Notably, CIC-dependent enrichment of BRG1/SIN3A/HDAC2 was not restricted to the *Vgf* promoter. Analysis of *Etv4* and *Etv5*, known CIC targets, also showed co-enrichment of CIC, BRG1, SIN3A, and HDAC2 within their promoter regions in control NPC (Fig 7C and 7D). Similar to the *Vgf* promoter, depletion of CIC reduced BRG1, SIN3A and HDAC2 occupancy within *Etv4* and *Etv5* promoter regions, which led to increased H3K27Ac and mRNA expression. Concordantly, depletion of CIC or BRG1 was sufficient to upregulate the mRNA expression of *Etv4* and *Etv5* in differentiating NPC (Fig 7E and 7F). Together, our results support a model that CIC functions as a neurogenic regulator by recruiting mSWI/SNF and SIN3-HDAC repressor complexes to transcriptionally regulate its target gene expression during neurogenesis (Fig 7G).

**Figure 7.**
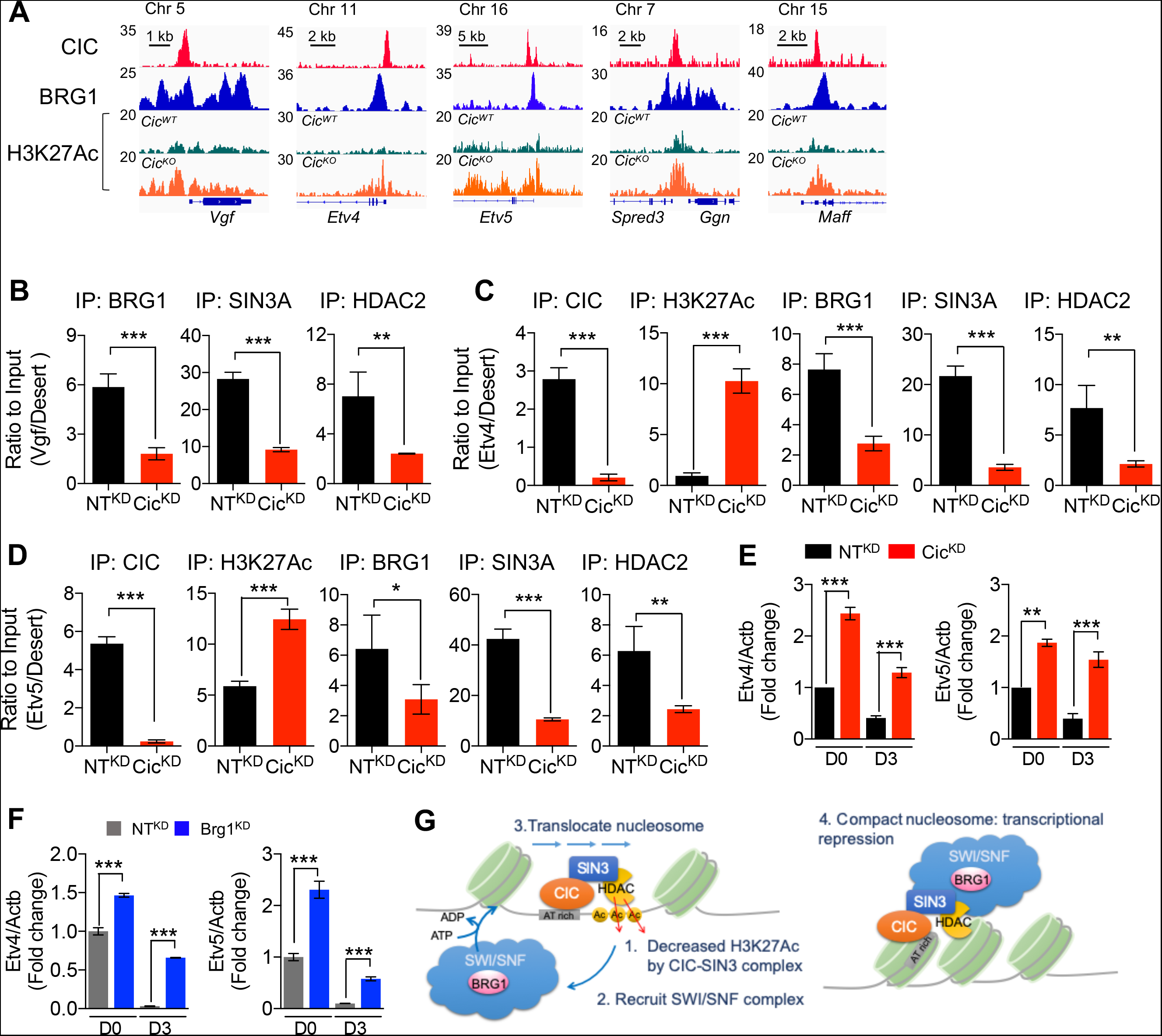
CIC represses VGF through tethering mSWI/SNF-SIN3 complex. A. ChIP tracks on CIC transcription targets. B-D. ChIP-qPCR results from NT^KD^ NPC and Cic^KD^ NPC at *Vgf* (B), *Etv4* (C), or *Etv4* (D) promoter region. Mean ± SEM of 3 experiments, **p<0.01, ***p<0.001 (Unpaired t-test). E. qPCR results of *Etv4* and *Etv5* mRNA expressions in NT^KD^ NPC and Cic^KD^ NPC at day 0 (D0) and day 3 (D3) of differentiation. Mean ± SEM of 3 experiments, **p<0.01, ***p<0.001 (One-way ANOVA). F. qPCR results of *Etv4* and *Etv5* mRNA expressions in NT^KD^ NPC and Brg1^KD^ NPC. Mean ± SEM of 3 experiments, *** p<0.001 (One-way ANOVA). G. The model schema for transcriptional regulatory mechanism by CIC-mSWI/SNF-SIN3 complex.

## Discussion

CIC is frequently mutated in oligodendrogliomas (Brat Verhaak et al., 2015). This compelling genetic evidence argues for its role in neuro-oncogenesis. But its function in brain development and tumorigenesis remains poorly understood. Recent studies reported smaller cerebral size and the behavioral deficit of forebrain targeted *Emx1-cre:Cic* knockout mouse (Lu et al., 2017). These and the current study commonly found the reduction in number of neurons in cortical layer 2-4. Another study using *Foxg1-cre* driven *Cic* knockout animals showed increased glia at the expense of neuronal differentiation (Ahmad et al., 2019). Considering the impaired neuronal maturation or terminal neural cell differentiation may underlie neuro-oncogenesis, we hypothesized that CIC functions in neuronal differentiation and development.

The reduced cortical thickness and reduced hippocampal dentate gyrus neuronal density phenotype of broadly deleting *Nes-cre* driven *Cic* knockout brain was consistent with that of forebrain-specific *Emx1-cre* driven *Cic* knockout animals (Lu et al., 2017). We suspect loss of CIC during embryonic brain development impaired neuronal differentiation and maturation as *Cic* null P0.5 brain already showed decreased neuron specific gene expression (Fig EV2A). This is consistent with the report by Ahmad et al., (Ahmad et al., 2019). We also observed that reduced overall brain size became progressively pronounced postnatally, a finding consistent with reported phenotype of *Foxg1-cre:Cic^L/L^* animals. The newborn brain continues to grow by a rapid increase in number of axons, dendrites, synapses, and glial cells. Thus, postnatal brain growth problems of CIC deficient brains may arise from disrupted connectivity such as defective elaboration of axons and dendrites, spinogenesis and maturation, synaptogenesis and remodeling, gliogenesis, and myelination. We speculate that decreased maturation of neurons contribute to the phenotype as the loss of CIC expression has a profound impact on the expression of dendrite and synaptogenesis genes based on RNA-seq analysis (Fig 1E). Gliogenesis and myelination changes appear less likely causes. *Emx1-cre:Cic* (Lu et al., 2017) or *Nes-cre:Cic* knockout mice showed normal range of corpus callosum thickness (Fig EV2D). In contrast, previous studies emphasized altered neural stem cell fate with expansion of OPC in CIC deficient mouse brains. *Foxg1-cre:Cic* knockout animals at P2 displayed decreased corpus callosum and *Mbp1/Cnp* levels due to defective terminal differentiation of OPC (Ahmad et al., 2019). Similarly, increased OLIG2-positive OPC in whole body *Cic* knockout at P28 and decreased oligodendrocytes differentiation of *Cic* knockout neural stem cells were noted (Yang et al., 2017). In our study, however, oligodendrocytes (OLIG2+), OPC (PDGFR +), and astrocytes (GFAP+) were not significantly expanded in broadly *Cic* deleted postnatal brains. Also, MBP staining did not show appreciable differences suggesting unaltered terminal differentiation of OPC into myelinating oligodendrocytes (Fig EV2D). These phenotypes suggest glial cell development was only marginally impacted in CIC null brain.

Notably, *Nes-cre* driven *Cic* knockout animals showed decreased postnatal growth leading to mortality around weaning age. This is consistent with the lethality of previously reported telencephalon-targeted *Foxg1-cre* driven *Cic* knockout animals (Ahmad et al., 2019). Based on the spectrum of different cre drivers it is likely that *Cic* deletion in the basal ganglia may be the cause of lethality. The physiological role of CIC in keeping the viability of animals warrants further investigation.

Previous studies demonstrated that VGF-derived peptide, TLQP-62, the C-terminal 62 amino acid peptide enhances neurogenesis of hippocampal cells *in vitro* and *in vivo* (Thakker-Varia, Behnke et al., 2014). Interestingly, VGF acted on early phases of neurogenesis by promoting neural progenitor proliferation while it inhibited terminal neuronal differentiation. It is noteworthy that CIC expression is dynamically upregulated through neuronal differentiation (Fig 2A, 2C, and EV3A). In the hippocampal dentate gyrus neurogenesis, where CIC has clear impact, we found that nuclear CIC is readily detected beginning in immature neurons with low TBR1 expression. The role of CIC lies at the transition from high TBR1 to low TBR1 as suggested by the expansion of high TBR1 cells upon the loss of CIC. These findings are concordant with the mechanism connecting CIC to VGF repression during neurogenic differentiation.

Under pathological condition of malignant gliomas, the neurotrophin BDNF induces glioma stem cells to secrete the VGF that promotes cancer cell growth and self-renewal in both autocrine and paracrine manner (Wang et al., 2018). Considering pro-tumorigenic action of VGF and its aberrant expression upon the deletion of *Cic* we hypothesized that it may promote hypercellularity or tumorigenic phenotypes in *Cic* knockout brains. To circumvent the lethality we induced conditional deletion of *Cic* before P10 in *Nestin-Cre^ERT2^:Cic^L/L^* by tamoxifen injection. Notably, we did not observe overt histological abnormality or tumor formation even with the introduction of *IDH1^R132H^* and *PIK3CA^E545K^* mutant alleles (unpublished observation). These findings argue for the insufficiency of these mutations for oligodendrogliomagenesis. It is conceivable that additional loss of 1p-located FUBP1 may facilitate tumorigenesis based on our previous study demonstrating its tumor suppressor role by promoting terminal neuronal differentiation (Hwang et al., 2018).

In our study, by combined ChIP-seq and RNA-seq analysis of P0.5 brains we found previously identified bona fide transcriptional targets of CIC including negative regulators of RTK/RAS/MAPK pathway (e.g., *Dusp4, Spred3, Spry4*, and *Nf1*) and the PEA3 subfamily of ETS transcription factors (*e.g., Etv4, Etv5*). These were all upregulated in *Cic* deficient brains. Those repressed targets showed CIC peak localized near TSS and largely overlapping with BRG1 agreeing with their molecular interaction. These CIC peaks co-occurred with H3K27Ac peaks whose coverage is conspicuously increased upon the loss of CIC expression. This finding is consistent with previous study demonstrating SIN3-HDAC corepressor complex recruitment to target loci by CIC. Indeed, we found decreased SIN3A and HDAC2 on target promoters in *Cic* knocked-down cells supporting our model (Fig 7G).

Interestingly, a subset of genes with intronic CIC peaks (*e.g., Satb2, Mef2c*, and *Myt1l*) showed downregulation in *Cic* knockout brains. These peaks, however, did not colocalize with H3K27Ac or BRG1 peaks unlike genes that are repressed by CIC (Appendix Fig S2). This lack of overlap excludes its role on enhancer or promoter-dependent regulation of those genes. Professional lineage determining transcription factors like OLIG2 showed distinct mechanism of target gene regulation engaging BRG1 containing SWI/SNF complex. It pre-patterns the recruitment of SWI/SNF complex to the enhancer of oligodendrocyte-specific genes (Yu et al., 2013). SWI/SNF complex recruited to distal lineage specific enhancers interacts with p300 to modulate histone H3 lysine 27 acetylation (Alver, Kim et al., 2017). In contrast, CIC-mediated transcriptional regulations primarily occurred at the proximal to the TSS unlike those lineage-determining transcription factors. This is likely due to the acute and dynamic regulation of CIC activity by RAS/MAPK signaling pathway (Weissmann et al., 2018). CIC integrates extracellular signals to maintain the homeostasis. Its role is likely to facilitate the transition of differentiating cells by coordinating gene expression according to the extracellular cue. Whether and how intronic CIC peaks regulate the gene expression warrants further investigation.

In conclusion, these molecular mechanisms outlining how CIC regulates neuronal differentiation can help our understanding of CIC-deregulated gliomagenesis and facilitate the development of novel therapeutic strategies.

## Materials and Methods

### Generation of *Cic^L/L^* Mouse

All animal use was approved by the Institutional Animal Care and Use Committee of the Weill Cornell Medicine. Mice were maintained on a 12 h light/dark cycle, and food and water was provided ad libitum. All mice were healthy with no obvious behavioral phenotypes, and none of the experimental mice were immune compromised. For all mouse studies, mice of either sex were used and mice were randomly allocated to experimental groups. Animals aged from postnatal day P0.5 to postnatal day P14 were used. Specific end-point developmental ages used for each experiment are indicated in the figure legends. *Cic* targeting vector was acquired from the KOMP repository of knockout mouse project (Vector # PG00139_Y_4_H08 – Cic, KOMP). The vector element is depicted in Fig EV1A. 144 ES cells clones were screened for homologous recombination. Two clones well targeted injected into blastocysts. A clone generated high percentage chimeras and achieved germ-line transmission when crossed to C57BL/6 females. Heterozygous *Cic^neo-loxP/+^* mouse was crossed with *FLPe* transgenic mouse (JAX #003800) for recombination of Frt. *Cic^L/L^* mice was crossed to *Nes-cre* transgenic mouse (JAX #003771) for recombination of *loxP* sites. The recombination was confirmed by PCR. Mouse genotypes were determined by PCR using toe DNA.

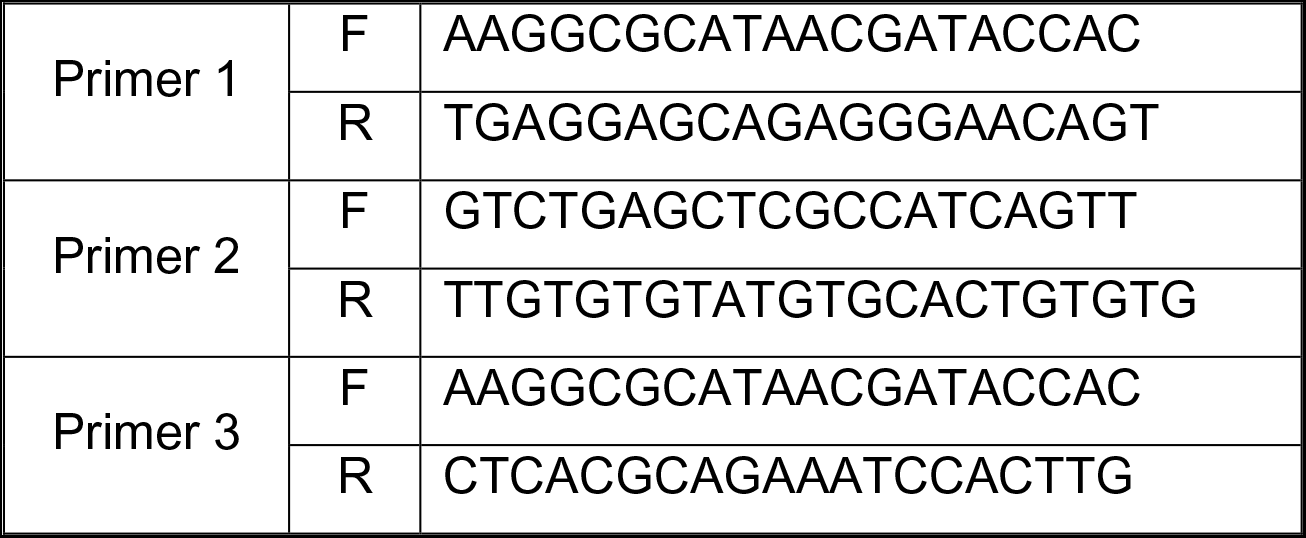

### NPC Cultures and Differentiation into Terminally Differentiated Neurons

*Ink/Arf*^-/-^ NPC (Khoo, Carrasco et al., 2007) were transduced with viral particles of pInducer-ASCL1 (NPC). pDONR221-ASCL1 (DNAsu #HsCD0004025) was gateway cloned into pInducer vector (Addgene #44012). After selecting the culture with hygromycin 50 μg/ml (Millipore Sigma #31282-04-9), the cells were divided and transduced with virus encoding shRNA for *Cic* (Millipore Sigma #TRCN0000302060 and #TRCN0000082010), non-Target shRNA for control (Millipore Sigma #SHC016), pLu-VGF-Flag (Homemade), or lentiCrisprV2-sgRNA (Addgene #52961). After selected with puromycin (3 μg/ml, InvivoGen # anti-pr-1) or blasticidin (10 ug/ml, InvivoGen # anti-bl-1), the all NPC are seeded on plate coated with fibronectin and poly-L-ornitine. On day 0, culture medium was replaced with N2 containing BDNF (10 ng/mL, PeproTech #450-02), NT-3 (10 ng/mL, PeproTech #450-03), B27 (ThermoFisher Scientific #1704044), and/or doxycycline (2 μg/mL, Research Products Int Corp. #D43020). On day 2, 0.5 % FBS was added to the medium to support astrocyte survival and the medium was changed every 2 days.

### Productions of Viruses

1.5 x 10^7^ 293T cells were seeded in 150 mm tissue culture dishes. After 24 h, the medium was replaced and cells were transfected with 4.5 μg of pMD2.G (Addgene #12259), 9 μg of psPAX2 (Addgene #12260), and 18 μg of target plasmid. After 48 h and 72 h of transfection, the medium containing viral particles was collected and was cleaned by filtration through a 0.45 μm cellulose acetate membrane. The viral particles were concentrated by ultracentrifugation for 2 h at 23,000 rpm and then, viral pellet was resuspended with 1 ml of Opti-MEM overnight at 4°C and stored in aliquots at −80°C.

### Generation of Knockdown Cells with CRISPR-Cas9 Strategy

sgRNAs were cloned into lentiCrisprV2 as previously published (Sanjana, Shalem et al., 2014). Viral particles were produced and infected to NPC followed by puromycin (3 μg/ml) selection. Gene targeting was confirmed by IF and WB analysis. Target sequences for sgRNAs are listed in following table.

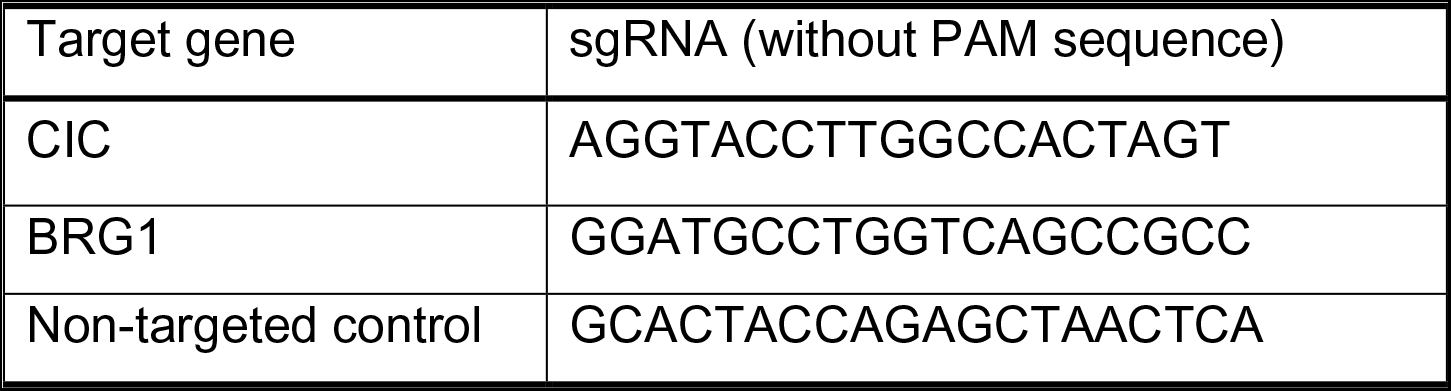

### Immunofluorescence Analysis

Cells were fixed with 4 % paraformaldehyde for 15 min at room temperature followed by permeabilization with 0.2 % triton X-100 in PBS. The cells were subjected to immunofluorescence staining with anti-CIC (1:500, (Lam et al., 2006)), anti-SATB2 (1:100, Abcam #ab51502), anti-CTIP2 (1:300, Abcam #ab18465), anti-BRG1 (1:1,000, Millipore Sigma #07-478), VGF (1:300, Abcam # ab69989), anti-SVIL (1:1,000, Millipore Sigma #S8695), anti-Nestin (1:1,000, DSHB # Rat-401), anti-GFAP (1:2,00, Origene # UM5000005), anti-NeuN (1:300, Cell signaling technology #24307), anti-MAP2a/b (1:100, Abcam #36447), anti-DCX (1:300, SantaCruz #SC-8066), anti-TBR1 (1:100, Abcam #ab31940), anti-TBR2 (1:100, Abcam #ab23345), or anti-SOX2 (1:500, Abcam #ab97959) antibodies incubated overnight at 4 °C and followed by labeling with Alexa 488-labeled anti-rabbit and Alexa 568-labeled anti-mouse secondary antibodies at room temperature for 1 h. For BrdU immunofluorescence staining, cells were incubated with 0.5 μg/ml of BrdU (Millipore Sigma #B5002) for 3 h followed by 1N HCl for 10 min on ice and 2 N HCl at 37 °C 20 min and neutralization with 0.1 M sodium borate buffer pH 8.5 for 10 min at room temperature. Immunofluorescence staining was performed with anti-BrdU (1:500, Dako #M0744) antibody overnight at 4°C followed by labeling with Alexa 568-labeled anti-mouse secondary antibody at room temperature for 1 h. Images were acquired using EVOS FL Cell Auto imaging system (ThermoFisher Scientific #AMAFD1000).

### Western Blot Analysis

Cells were lysed by RIPA buffer or Laemmli buffer followed by sonication (30 watt/5 sec/10 cycles). Protein concentration was determined by using Pierce^TM^BCA protein assay kit. 30 μg of proteins were fractionated by SDS-PAGE electrophoresis and transferred to PVDF membrane (ThermoFisher Scientific #88518) using a transfer apparatus followed by manufacturer’s instructions. After incubation with 5 % skim milk in TBST (10 mM Tris, pH 8.0, 150 mM NaCl, 0.5 % Tween 20) for 1 h, the membrane was incubated with antibodies against β-actin, CIC (1:1,000), SATB2 (1:500), CTIP2 (1:2,000), NeuN (1:1,000), DCX (1:2,000, Cell signaling technology #4604), BRG1 (1:3,000), SVIL (1:1,000), α Tubulin (1:10,000, clone4A1, DSHB #AB_2732839), OLIG2 (1:5,000, Millipore Sigma #AB9610), PDGFRA (1:1.000, Cell signaling technology #3174), CNPase (1:2,000, Abcam #ab6319), GFAP (1:5,000), or VGF (1:2,000) overnight at 4 °C. Membranes were washed three times with TBST for 30 minutes and then incubated with HRP conjugated anti-mouse, anti-rabbit, or anti-guinea pig (1:10,000) diluted in 3 % skim milk for 1 h. Blots were washed with TBST three times and developed with the SuperSignal™ West Pico Chemiluminescent substrate (ThermoFisher Scientific #34080) according to the manufacturer’s protocols.

### Immunoprecipitation Analysis

Cell or tissues were lysed using 1 % TNT buffer (135 mM NaCl, 20 mM Tris-HCl, 1 mM EDTA, and 1% Triton X-100) for 30 min on ice. After centrifugation to remove the debris, 500 μg of protein was incubated with 1-3 μg of antibody at 4 °C for 16 h. 15 μl of Dynabeads® Protein G (ThermoFisher Scientific #1009D) was added and incubated 4 °C for 3 h. The beads were washed 3 times with 1 % TNT buffer and eluted the proteins with 2X Laemmli buffer. The protein interaction was determined by western blot protocol.

### Quantitative Real-time PCR (qPCR) Analysis

Total RNAs were extracted from cells or tissues by using NucleoSpin RNA kit (Macherey-Nagel #740955.25). Reverse transcription was carried out on 500 ng of total RNA using utilizing RevertAid RT kit (ThermoFisher Scientific #K1691). qRT-PCR was performed on cDNA samples using the PowerUp™ SYBR® Green Master Mix (ThermoFisher Scientific #A25778) and was performed the qPCR on the 7500 Fast Real-time PCR system (ThermoFisher Scientific #4351106). Primer sequences are as below. Each sample was run as duplicates and the mRNA level of each sample normalized to that of ACTB mRNA. The relative mRNA level was presented as unit values of 2^dCt (=Ct of ACTB-Ct of gene).

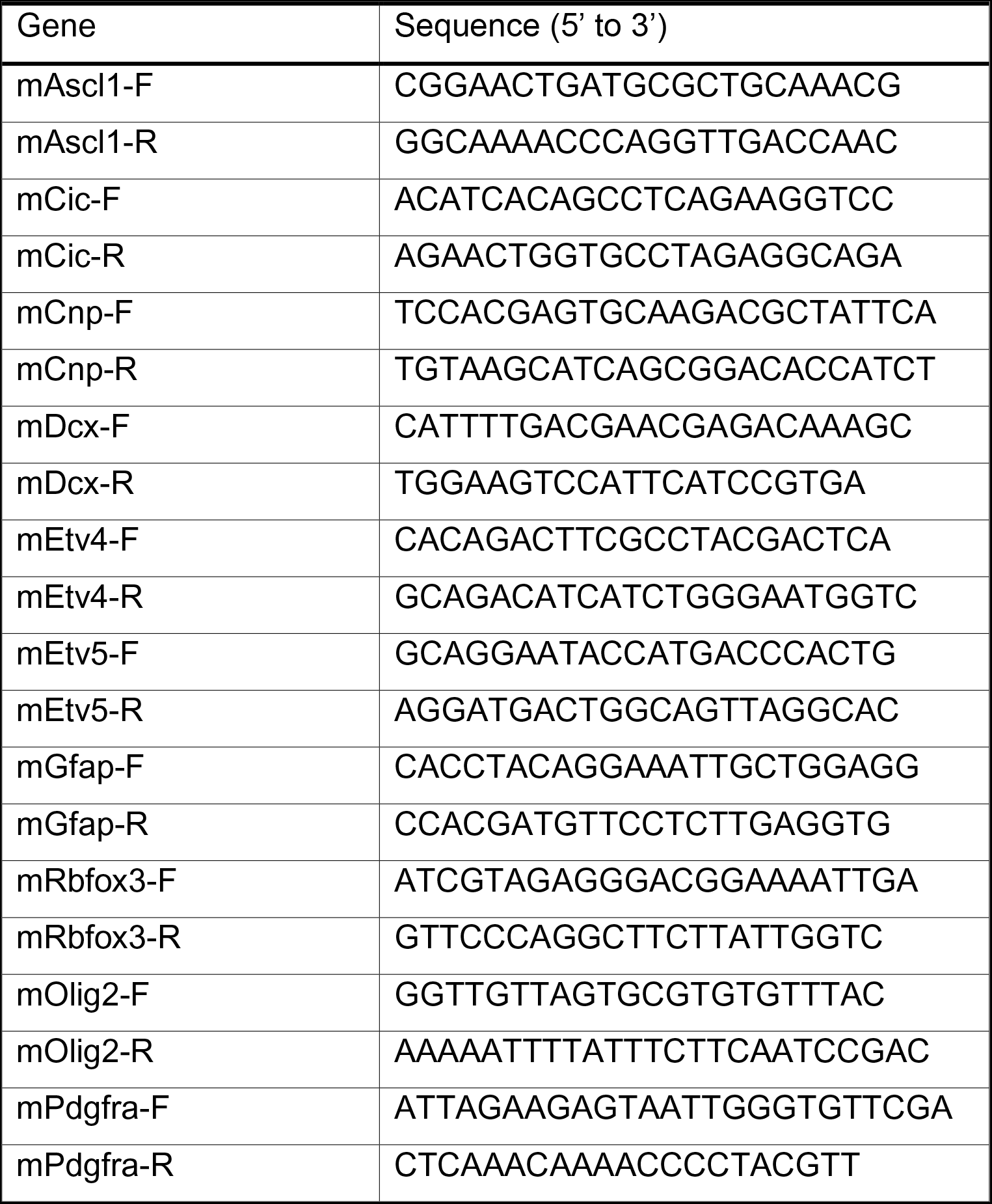
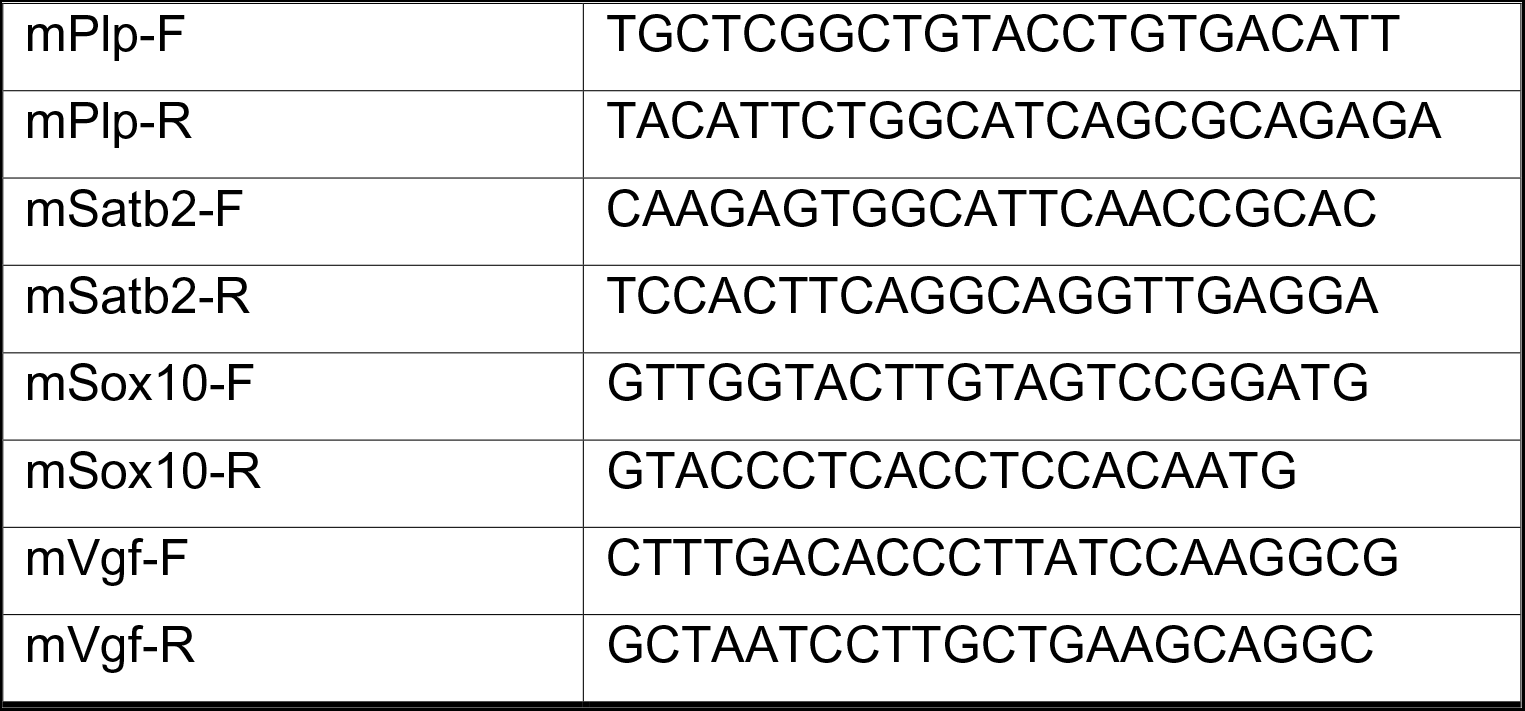

### Mass Spectrometry Analysis

The analysis was performed at the Proteomics and Metabolomics Core facility of Weill Cornell Medicine. In-gel digestion was performed according to a previous published protocol (Shevchenko, Wilm et al., 1996). Briefly, gel pieces were excised and distained, followed by reduction, alkylation and digestion with trypsin. The peptides were then extracted from the gels, desalted and analyzed by liquid chromatography-tandem mass spectrometry (LC-MS/MS). An EASY-nLC 1200 coupled on-line to a Fusion Lumos mass spectrometer was used for LC-MS/MS analysis. Buffer A (0.1 % formic acid in water) and buffer B (0.1 % formic acid and 80 % acetonitrile in water) were used as mobile phases for gradient separation. A 75 µm x 15 cm chromatography column was packed in-house for peptide separation. Peptides were separated with a gradient of 5–40 % buffer B over 20 min, 40-100% B over 5 min at a flow rate of 300 nL/min. The Fusion Lumos mass spectrometer was operated in data dependent mode with 1 s cycle time. Survey scans were acquired in the Orbitrap mass analyzer over a range of 300-1500 m/z with resolution 120,000 at m/z 200. The most abundant precursors from the survey scan were selected with an isolation window of 1.6 Thomsons and fragmented by higher-energy collisional dissociation with normalized collision energy of 35. MS/MS scans were acquired in the ion trap mass analyzer with rapid scan rate. The automatic gain control target value was 1e6 for MS scans and 1e4 for MS/MS scans respectively, and the maximum ion injection time was 60 ms for both. The raw files were processed using the MaxQuant computational proteomics platform (Cox & Mann, 2008)(version 1.5.5.1). The fragmentation spectra were searched against the UniProt mouse protein database (contain 80,593 sequences), and allowed up to two missed tryptic cleavages. Oxidation of methionine and protein N-terminal acetylation were used as variable modifications for database searching. Carbamidomethylation of cysteine was used as a fixed modification. The precursor and fragment mass tolerances were set to 7 ppm and 0.5 Da, respectively. Both peptide and protein identifications were filtered at 1 % false discovery rate (FDR).

### RNA-seq and Data Analysis

Total RNAs were isolated from P0.5 brains of *Cic^WT^* and *Cic^KO^* mouse and subjected to RNA sequencing at the Genomics Resources Core facility of Weill Cornell Medicine. RNA-seq libraries were prepared using the Illumina TruSeq stranded mRNA library preparation kit and sequenced on HiSeq4000 sequencer (Illumina). RNA-seq data were aligned to the mm9 reference genome using STAR 2.3.0e (Dobin, Davis et al., 2013). Raw counts of each transcript were measured by featureCounts v1.4.6-p5 (Liao, Smyth et al., 2014). Lists of differentially expressed genes were generated by DESeq2-1.4.5 in R (Love, Huber et al., 2014). GSEA analysis in this manuscript were generated from GSEA preranked model. The input of GSEA analysis is the gene expression level logFC (*Cic*^KO^ mouse versus control). Pathway analysis (Fig 3E) was performed using DAVID 6.8 (Huang, Sherman et al., 2009a, Huang, Sherman et al., 2009b).

### ChIP-Seq and qPCR analysis

We used whole brain from P0.5 postnatal mouse or 10^7^ of neural progenitor cells for each analysis. ChIP analysis was performed with modifications (Lee, Johnstone et al., 2006). In brief, tissues or cells were reacted with 5 mM EGS (ThermoFisher Scientific #21565) for 45 minutes at room temperature to cross-link protein-protein interaction. After centrifugation and removal of supernatant, cross-linked for 10 minutes with 1% paraformaldehyde and quenched with 120 mM glycine for 5 minutes at room temperature. After nucleus isolation, the chromatin was digested with 6,000 gel units of micrococcal nuclease for 10 minutes at 37°C. The enzyme reaction was quenched by adding 20 mM EDTA. Resuspended the pellet in shearing buffer (50 mM Tris-HCl, 10 mM EDTA, and 0.1 % SDS) and broke the nuclear membrane using the Covaris M220 Focused-ultrasonicator according to the manufacturer’s instructions. Immunoprecipitation was performed with 10 μg of anti-CIC, anti-BRG1 (Cell signaling technology #49360), anti-H3K27Ac (Abcam #4729), anti-Sin3A (Abcam #ab3479), or anti-HDAC2 (Cell signaling technology #57165) overnight at 4 °C. 30 μl of pre-cleared Dynabeads® Protein G was added and incubated for 3 h at 4 °C. Washed the beads by high salt buffer (50 mM HEPES, 500 mM NaCl, 1mM EDTA, 0.1 % SDS, 1 % Triton X-100, and 0.1 % sodium deoxycholate) and RIPA buffer (including LiCl) and eluted the chromatin by elution buffer (50 mM Tris-HCl, 10 mM EDTA, and 1% SDS). After treated with RNase and Proteinase K, the DNA was incubated at 65 °C overnight to reverse cross-linking. DNA was extracted using NucleoSpin Gel and PCR clean-up DNA extraction kit (Macherey-Nagel #740609.25) and carried out size-selection to obtain ∼300 bp of DNA fragments using SPRIselect Reagent (Beckman Coulter # B23317). RT-qPCR was performed using specific primers described in the following table.

Libraries were made using KAPA Hyper Prep kit (KAPA Biosystems # KR0961) following manufacturer’s instructions. Briefly, 30 ng from each immunoprecipitated or Input DNA, were end-repaired, phosphorylated, A-tailed and ligated to adaptors. Ligated products were size selected with 0.8X SPRI beads to obtain 250∼350 bp of DNA. After purification, 8-cycle PCR amplification reaction was performed. PCR product was cleaned by the use of 1X SPRI beads. Final product was resuspended in 30 μl of Tris buffer. Final yields were quantified in a Qubit 4.0 Fluorometer and quality of the library was assessed by running on a DNA1000 Bioanalyzer chip. Libraries were normalized to 2 nM and pooled at the desired plexity. Sequencing and post-processing of the raw data was performed at the Epigenomics Core at Weill Cornell Medicine as follows: Libraries were clustered at 6 pM on single read flow cell and sequenced for 50 cycles on an Illumina HiSeq 2500. Illumina’s CASAVA 1.8.2 software was used to perform image capture, base calling and demultiplexing.

ChIP-seq data were aligned to the mm9 reference genomes using bowtie-0.12.9 with default paramters -n 2 and -best (Langmead, Trapnell et al., 2009). Peak calling was performed by macs14 1.4.2 (Zhang, Liu et al., 2008) with default parameters. BigWig files of ChIP-seq data track and analysis of read density in peak regions were generated using deeptools 3.1.3 (Ramírez, Ryan et al., 2016). Read density of specific genomic regions were displayed using Integrative Genomics Viewer (IGV) 2.4.19 (Robinson, Thorvaldsdóttir et al., 2011).

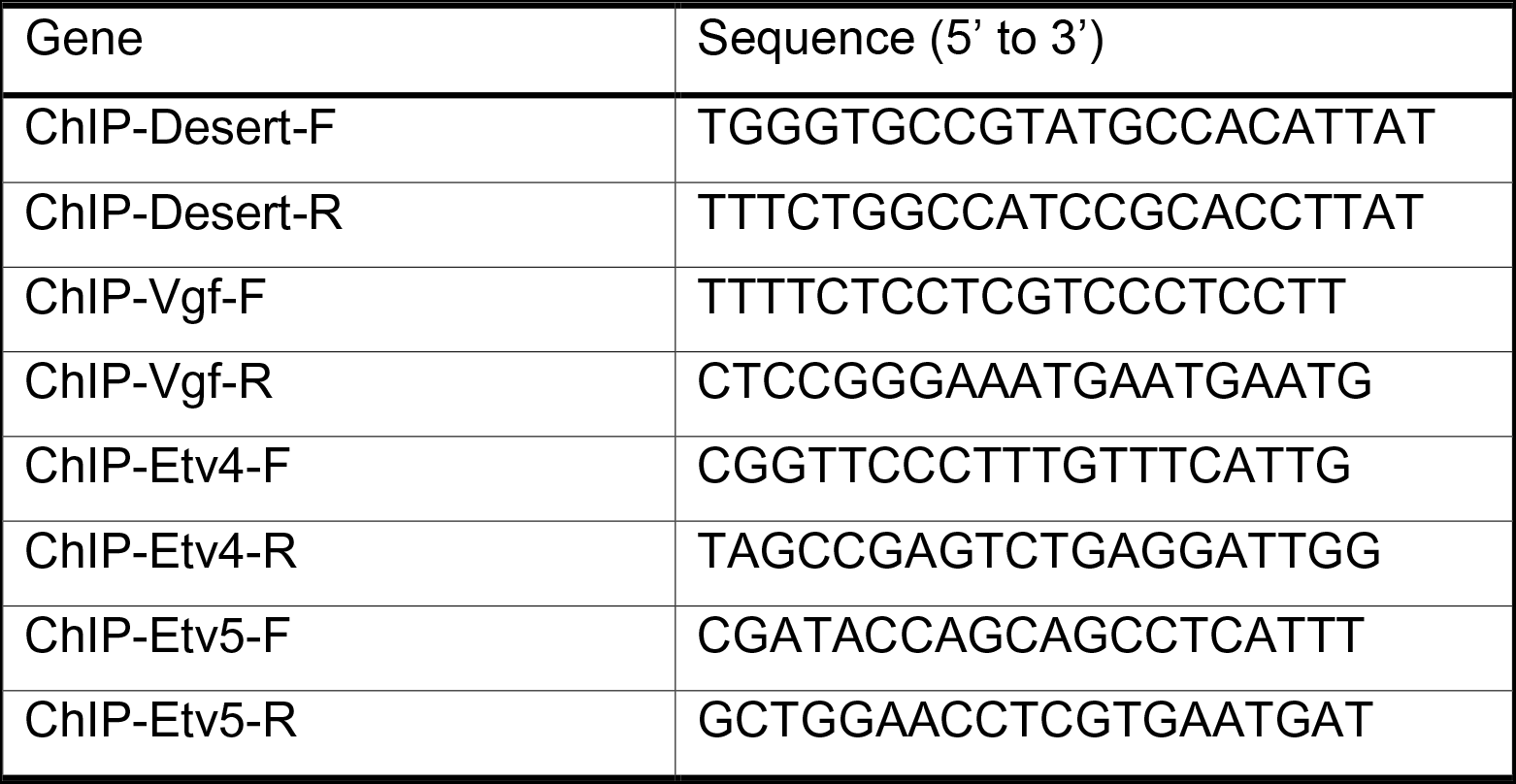

### Quantification and Statistical Analysis

We determined experimental sample sizes on the basis of preliminary data. All results are expressed as mean ± SEM. GraphPad Prism software was used for all statistical analysis. Normal distribution of the sample sets was determined before applying unpaired Student’s two-tailed t-test for two group comparisons. One-way ANOVA was used to assess the differences between multiple groups. The mean values of each group were compared by the Bonferroni’s post-hoc procedure. Differences were considered significant when P<0.05.

## Data Availability

The datasets and computer code produced in this study are available in the following databases:

- RNA-seq and ChIP-seq data: Gene expression NCBI GEO: GSE131302 (https://www.ncbi.nlm.nih.gov/geo/query/acc.cgi?acc=GSE131302)

## Acknowledgements

Authors thank Dr. Yoontae Lee for providing CIC expression plasmids and Dr. Huda Zhoghbi for CIC antisera which was used for the initial optimization of CIC immunoblotting. We thank Dr. Zhe Cheng and Dr. Guoan Zhang at the Proteomics and Metabolomics Core Facility and Dr. Tuo Zhang at the Genomics Core Facility of The Weill Cornell Medicine for mass spectrometry and sequencing data analysis. J.P. is supported by the Irma T. Hirschl Award, and National Institutes of Health Grant AG048284 and CA214274.

## Author Contributions

Conception and design: I.H., J.P.

Development of methodology: I.H., H.P.

Acquisition of data: I.H.

Analysis and interpretation of data (e.g., statistical analysis, biostatistics, computational analysis): I.H., J.Y., H.P., H.Z., J.P.

Writing, review, and/or revision of the manuscript: I.H., H.Z., J.P.

Administrative, technical, or material support (i.e., reporting or organizing data, constructing databases): I.H., O. E., H.Z., J.P.

Study supervision: J.P., O.E.

